# Changes of oscillatory and aperiodic neuronal activity in working memory following anaesthesia: a prospective observational study

**DOI:** 10.1101/2022.05.13.491765

**Authors:** Janna D. Lendner, Ulrich Harler, Jonathan Daume, Andreas K. Engel, Christian Zöllner, Till R. Schneider, Marlene Fischer

## Abstract

**Background:** Anaesthesia and surgery can lead to cognitive decline, especially in the elderly. However, to date, the neurophysiological underpinnings of perioperative cognitive decline remain unknown.

**Methods:** We included male patients, who were 60 years or older scheduled for elective radical prostatectomy under general anaesthesia. We obtained neuropsychological (NP) tests as well as a visual match-to-sample working memory (WM) task with concomitant 62-channel scalp electroencephalography (EEG) before and after surgery.

**Results:** A total number of 26 patients completed neuropsychological assessments and EEG pre- and postoperatively. Behavioural performance declined in the neuropsychological assessment after anaesthesia (total recall; *t-tests:* t_25_ = -3.25, Bonferroni-corrected p = 0.015 d = -0.902), while WM performance showed a dissociation between match and mis-match accuracy (*rmANOVA*: match*session F_1,25_ = 3.866, p = 0.060). Distinct EEG signatures tracked behavioural performance: Better performance in the NP assessment was correlated with an increase of non-oscillatory (aperiodic) activity, reflecting increased cortical activity (*cluster permutation tests:* total recall r = 0.66, p = 0.029, learning slope r = 0.66, p = 0.015), while WM accuracy was tracked by distinct temporally-structured oscillatory theta/alpha (7 – 9 Hz), low beta (14 – 18 Hz) and high beta/gamma (34 – 38 Hz) activity (*cluster permutation tests:* matches: p < 0.001, mis-matches: p = 0.022).

**Conclusions:** Oscillatory and non-oscillatory (aperiodic) activity in perioperative scalp EEG recordings track distinct features of perioperative cognition. Aperiodic activity provides a novel electrophysiological biomarker to identify patients at risk for developing perioperative neurocognitive decline.

## Introduction

Postoperative cognitive impairment after surgery and anaesthesia includes postoperative delirium (POD)^1^, delayed neurocognitive recovery (DNCR) and postoperative neurocognitive disorders^2, 3^, all of which are associated with increased morbidity and mortality.^1–3^ Elderly patients above 60 years are at high risk^2, 4, 5^ to suffer from perioperative cognitive decline with incidences of up to 25 %.^3^ Several cognitive domains such as attention, memory, and executive functions can be affected by DNCR.^6^ Working memory (WM) maintenance is one function that may be affected postoperatively even in the long term, which has been shown in both animal and human studies.^7–9^ Neurophysiological studies in healthy human participants showed specific patterns of oscillatory neuronal activity in several frequency bands during the maintenance interval of WM tasks.^10–13^ Therefore, this study aimed at investigating the changes of oscillatory activity before and after general anaesthesia for elective non-cardiac surgery in a visual WM task.

In addition to changes in oscillatory neuronal activity after surgery, we examined the modulation of aperiodic neuronal activity, which can be quantified by the spectral slope of the 1/f^x^ exponential function of the power spectrum (PSD).^14^ Recent evidence demonstrated that aperiodic activity contains rich information about the current behavioural state. It is related to arousal in sleep and anaesthesia,^15–18^ but also modulated during various cognitive tasks such as attention and working memory.^18, 19^

To address these outstanding questions, we recorded both a cognitive neuropsychological (NP) test battery, as well as a visual WM task with simultaneous whole-head 62-channel scalp EEG before and after general anaesthesia in patients undergoing radical prostatectomy to assess 1) perioperative cognitive performance and 2) concomitant alterations of electrophysiological patterns. Specifically, we hypothesized that anaesthesia alters oscillatory activity in the theta, alpha and beta band, as these frequencies showed pronounced changes during WM maintenance.^20^ In addition, we examined aperiodic activity, as measured by the spectral slope of the power spectrum^15–17^, across pre- and postoperative sessions and investigated how oscillatory and aperiodic electrophysiological signatures, relate to cognitive performance.

## Material and Methods

### Ethics approval

The study protocol was approved by the local ethics committee of the Hamburg Chamber of Physicians (protocol number PV4782; 27.04.2016). All study participants gave written informed consent before the initiation of study-related procedures.

### Design, setting and participants

This prospective observational study was performed at a high-volume prostate cancer centre in Northern Germany. Between June 2016 and July 2017, we included a convenience sample of patients aged 60 years or older, who spoke German fluently and were scheduled for robot-assisted radical prostatectomy. Patients with pre-existing cognitive impairment or cerebrovascular disease were excluded. Study participants completed a neuropsychological (NP) assessment for cognitive function and performed a WM task on the day before surgery *and* on one day post-surgery (2^nd^-4^th^ day). For details on data collection and anaesthesiologic management see Supplementary Information.

### Neuropsychological Assessment

The NP test battery included the California Verbal Learning Test (CVLT), the Trail Making Test (TMT), the Grooved Pegboard test (GPT) and the Digit Span Forward Test (DS; **Fig. 1**). To obtain standardized test scores for each subtest of the CVLT, we calculated the difference between pre- and postoperative scores for each patient and divided the result by the baseline standard deviation (SD). For the TMT, the GP and the DS z-scores were calculated accordingly after subtracting the practice effect. A negative z-score for CVLT total recall, CVLT learning slope as well as the digit span (and a positive z-score for TMT-B and DS) indicated a worse performance in the second compared to the first session.

**Figure 1:**
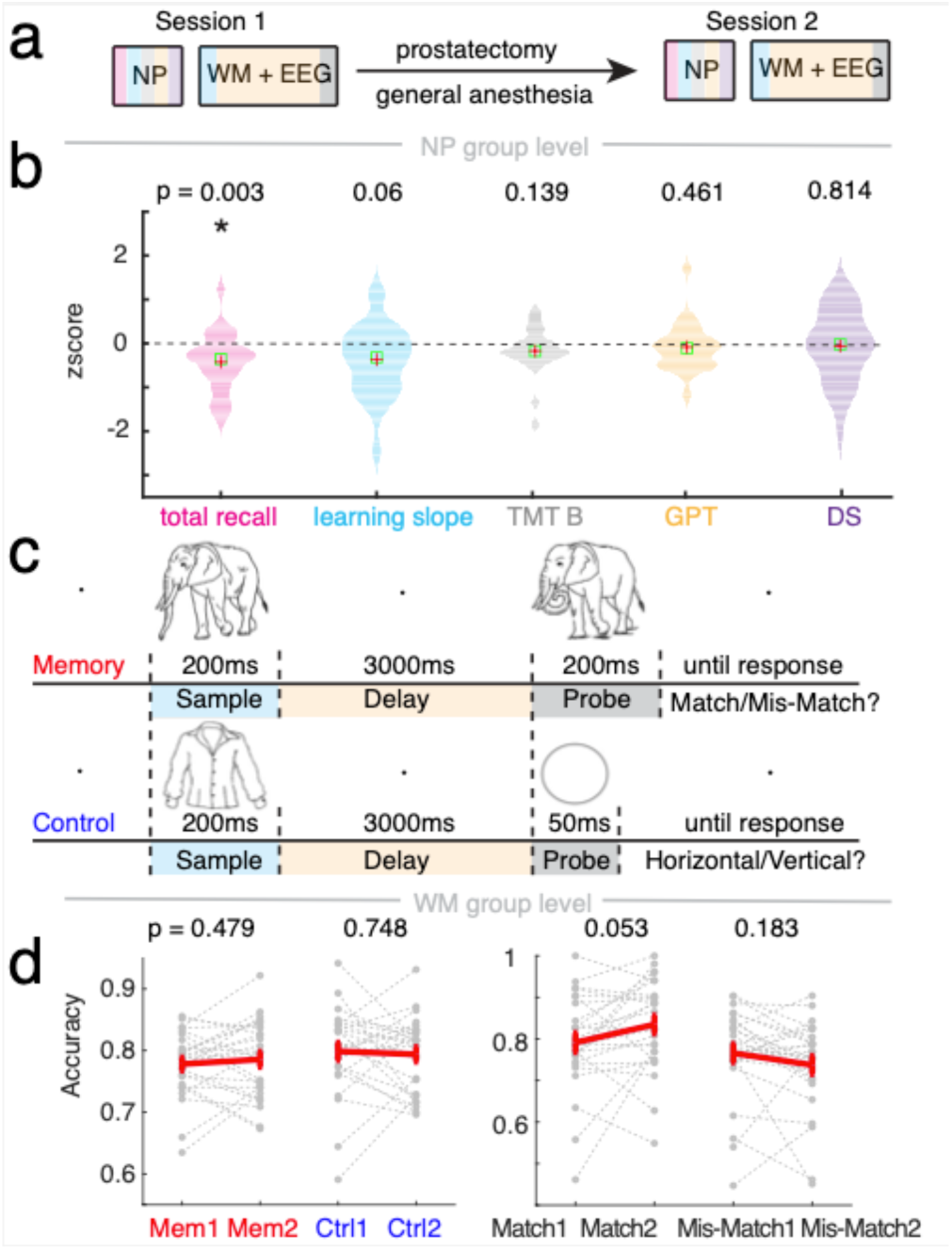
Cognitive performance before and after general anaesthesia. **a**, Study design: Neuropsychological (NP) tests and working memory (WM) task with electroencephalography (EEG) were obtained before (Session 1) and after (Session 2) prostatectomy under general anaesthesia. **b**, NP test scores (n = 26). Only California Verbal Learning Task total recall (pink) showed significant changes compared to zero (p-value corrected for multiple comparisons = 0.015; * p < 0.05), revealing that patients could remember less words. TMT-B (grey): Trail Making Test B, GPT (orange): Grooved Pegboard Test, DS (purple): Digit Span. Red cross: mean. Green box: standard error of the mean (SEM). **c**, Task design WM: Sample presentation (blue) is followed by 3 second delay (orange), then probe presentation (grey): in memory either exactly the same (match) or an altered version (non-match), in control, either a horizontally or vertically oriented ellipse (grey). **d**, Accuracy of memory and control (left) and match and mis-match memory trials (right; n = 26). Discrimination of matches was more accurate after anaesthesia while less mis-matches were identified (t-test, p-values uncorrected). Grey dotted lines: single subjects. Red lines: mean +/- SEM.

### Working Memory Task

For the assessment of task-related electrophysiologic changes, we chose a WM task, since impairment in WM has been implicated in perioperative cognitive disorders.^21–23^ Pre- and postoperative sessions consisted of two conditions – a visual delayed match-to-sample task (memory condition) as well as a non-mnemonic visual discrimination task (control condition) as described previously.^20^ Stimuli were black and white line drawings of natural objects (**Fig. 1**).^24^ In a total of 8 blocks (4 of each memory and control) with 52 trials each, patients completed 416 trials (208 of each condition) consisting of unique sample/probe pairs (task details see Supplemental Information).^20^

### Behavioural data analysis

Reaction time (RT in seconds) and accuracy (AC between 0 and 1, where 1 equals 100%) were determined for every condition and session (session 1 – before, session 2 – after anaesthesia; condition – memory or control). Memory performance was further split into match (RT_Match_/AC_Match_) and non-match trials (RT_Non-Match_/AC_Non-Match_) and then averaged across trials. Performance differences between matches and non-matches were calculated by subtracting the second from the first session.

### Electroencephalography

EEG data was recorded during the working memory and the control task from an equidistant 62 electrode scalp montage montage (Easycap, Herrsching, Germany) with two additional electrooculography (EOG) channels below the eyes and referenced against the tip of the nose. EEG was recorded with a passband of 0.016-250 Hz and stored with a sampling rate of 1000 Hz using BrainAmp amplifiers (BrainProducts, Gilching, Germany) in a dimly lit, sound-attenuated recording room.

### Electrophysiological data analysis

Data analysis was performed using Matlab R2018b (The MathWorks, Inc., Natick, Massachusetts, United States) and the FieldTrip^25^ toolbox (version 20172908) as well as custom code. For pre-processing, electrophysiological data was notch-filtered to remove line noise in 0.1 Hz steps (bandstop filter; 49 to 51 and 99 to 101 Hz) as well as high- (0.1 Hz) and lowpass-filtered (100 Hz). Data was then averaged using a common median reference excluding the two EOG channels. Electrophysiological data was epoched into 5-second segments time-locked to sample onset (−1 to +4 s).

Trials containing jumps or strong artifacts were detected using FieldTrip^25^ (ft_rejectvisual) and rejected after visual inspection. Next, independent component analysis (fastica^26^) was used to remove eye-blinks, horizontal eye-movements, electrocardiogram artifacts or broadband noise based on the component spectrum, topography and time course.

### Spectral power analysis

Time-frequency decomposition was performed on epoched data using a sliding window Fast Fourier Transformation (*mtmconvol*^25^) between -0.5 to 3.5 seconds in 50 ms steps and in the frequency range from 1 to 40 Hz in 1 Hz steps using a 500 ms Hanning taper with 50 % overlap. Each trial was then z-scored to a bootstrapped baseline (−500 to 0 ms) using a permutation approach (pooled baseline of all trials per channel; 1000 iterations).

### Spectral slope analysis

Calculation of spectral slope was performed on the PSD (settings see above) between 30 to 40 Hz, in line with recent work.^15^ For each channel and trial, we obtained a first-degree polynomial fit to the power spectra in *log-log* space (*polyfit*.*m*, MATLAB and Curve Fitting Toolbox Release R2018b, The MathWorks, Inc., Natick, Massachusetts, United States).

### Statistical analysis

For statistical comparison of behavioural data, we employed dependent samples t-tests. All significant results were Bonferroni-corrected for multiple comparisons, if not stated otherwise. Effect size was calculated using Cohen’s d. To determine the influence of conditions, sessions and trials type on behaviour, we utilized Greenhouse-Geisser corrected 2-way repeated measures analysis of variance (RM-ANOVA). To assess the spatial extent of the observed effects in EEG, we calculated cluster-based permutation tests to correct for multiple comparisons as implemented in FieldTrip^25^ (Monte-Carlo method; maxsum criterion; 1000 iterations) reporting p- as well as the sum of t-values. A permutation distribution was obtained by randomly shuffling condition labels. Spatial clusters are formed by thresholding dependent samples t-tests between control and memory as well as memory between sessions at a p-value < 0.05 for the comparisons of ERP (**Fig. S5**), power (**Fig. 3**) and spectral slope (5000 iterations; **Fig. 5**). To assess the spatial extent of a correlation between WM/NP behaviour and power/slope differences between memory sessions in the delay period (0.3 to 3.1 s; **Fig. 4, 5**), we used rank correlations at a p-value of < 0.05 for the cluster test. To control for an influence of WM behaviour on the NP-slope association, we utilized a partial correlation that partialled accuracy out before computing the correlation. The results of this analysis should be regarded as hypothesis generating.

## Results

A total of 41 male patients (for patient characteristics see **Table 1, S2**) were included in this study (flow chart see **Fig. S1**). Data analysis focused on 26 patients that completed both neuropsychology (NP) and working memory (WM) sessions pre- and post-operation, of whom seven fulfilled definition of DNCR (26.92 %). Demographic and clinical data of study participants including, who did not complete the postoperative assessments, are presented in Table S2.

**Table 1.**
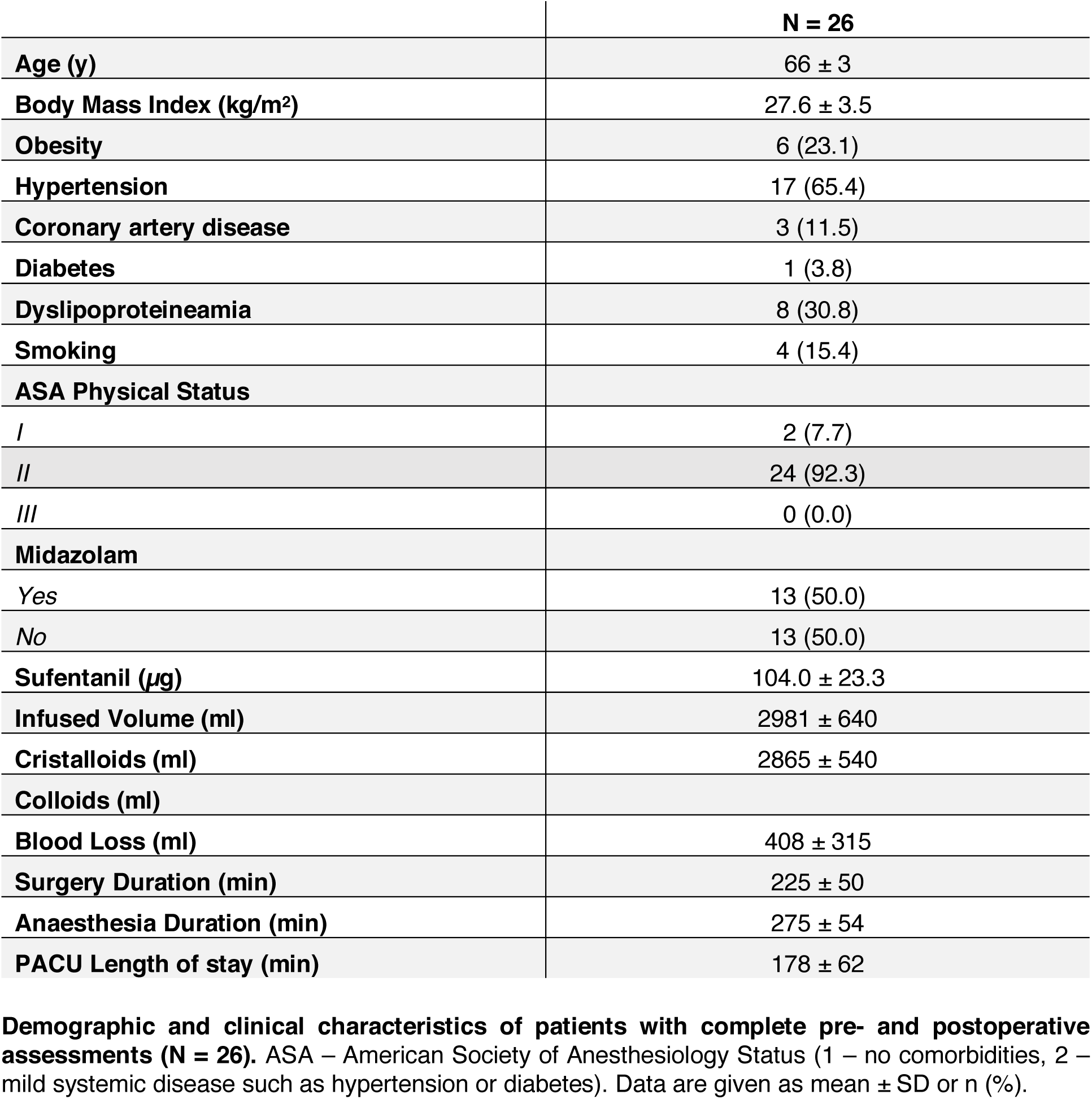

### Cognitive performance before and after general anaesthesia

Patients remembered less words in the post-anaesthesia NP session (CVLT total recall/zero t_25_ = -3.25, Bonferroni-corrected p = 0.015 d = -0.902; **Fig. 1b, Table S2**). In addition, they tended towards being slower in learning new words (CVLT learning slope/zero t_25_ = -1.97, Bonferroni-corrected p = 0.3, d = -0.546). The other NP tests did not differ between sessions (**Table S2, Fig. 1b**). Thus, subsequent analysis focused on CVLT metrics.

Between the two memory sessions of the WM task, overall accuracy was comparable (values see **Table 2** and **Fig. 1**; *repeated-measures ANOVA:* condition F_1,25_ = 4.574, p = 0.042; session F_1,25_ = 0.012, p = 0.912; condition*session F_1,25_ = 1.404, p = 0.247; dprime *t-test*: p = 0.245). When analysing match and mis-match trials separately, there was an overall tendency for patients to become more accurate in detecting matches and less so in recognizing mis-matches (**Fig. 1**; *repeated-measures ANOVA:* match F_1,25_ = 3.412, p = 0.077; session F_1,25_ = 0.497, p = 0.487; match*session F_1,25_ = 3.866, p = 0.060), in line with a response bias shift towards matches (**Table 2**; *t-test:* p = 0.049). For reaction times, there were no overall, match or mis-match differences (**Table 2, Fig. S2**), therefore, we focused on accuracy differences between match and mis-match trials for subsequent analysis of any association between WM behaviour and electrophysiology.

**Table 2.**

|  | Se 1 |  |  |  | Se 2 |  |  |  |
| --- | --- | --- | --- | --- | --- | --- | --- | --- |
|  | Mem | Ctrl | Match | Non-Match | Mem | Ctrl | Match | Non-Match |
| **Reaction Times (s)** | 1.104 ± 0.042 | 1.156 ± 0.081 | 1.131 ± 0.045 | 1.076 ± 0.044 | 1.119 ± 0.032 | 1.1484 ± 0.069 | 1.129 ± 0.037 | 1.109 ± 0.030 |
| **Accuracy** | 0.778 ± 0.011 | 0.7988 ± 0.014 | 0.791 ± 0.023 | 0.766 ± 0.022 | 0.786 ± 0.012 | 0.794 ± 0.012 | 0.835 ± 0.022 | 0.737 ± 0.022 |
| **Slope** | -0.272 ± 0.083 |  |  |  | -0.129 ± 0.098 |  |  |  |
| **D-prime** | 1.658 ± 0.074 |  |  |  | 1.758 ± 0.103 |  |  |  |
| **Bias** | -0.065 ± 0.074 |  |  |  | -0.213 ± 0.069 |  |  |  |

**Working memory performance (n = 26).** All values mean ± SEM. Se – session. Mem – memory, Ctrl – control condition.

Interestingly, there was no relationship between accuracy and NP assessment scores (Δ = Se2 – Se1; Δ ACmatch/ z-score total recall r = 0.17, p = 0.417; Δ ACmatch/z-score learning slope r = 0.26, p = 0.191; Δ ACmis-match/ z-score total recall r = 0.05, p = 0.817; Δ ACmis-match/z-score learning slope r = -0.07, p = 0.724; **Fig. S2a**) indicating that they reflect different cognitive processes.

### Oscillatory activity in the beta frequency band is associated with a visual accuracy

Grand average power (**Fig. 2**) across all subjects, conditions and sessions revealed an early increase of delta/theta power (0.5 – 7 Hz), followed by a decrease in the alpha/beta range (8 – 19 Hz) and an increment in the low and high beta frequency band (12 – 16, 20 – 26 Hz) compared to baseline (−0.5 – 0 s), in line with recent work^20^. Here, we focused on these frequency bands and the delay period (0.3 – 3.1 s) where stimulus related activity was absent.

**Figure 2:**
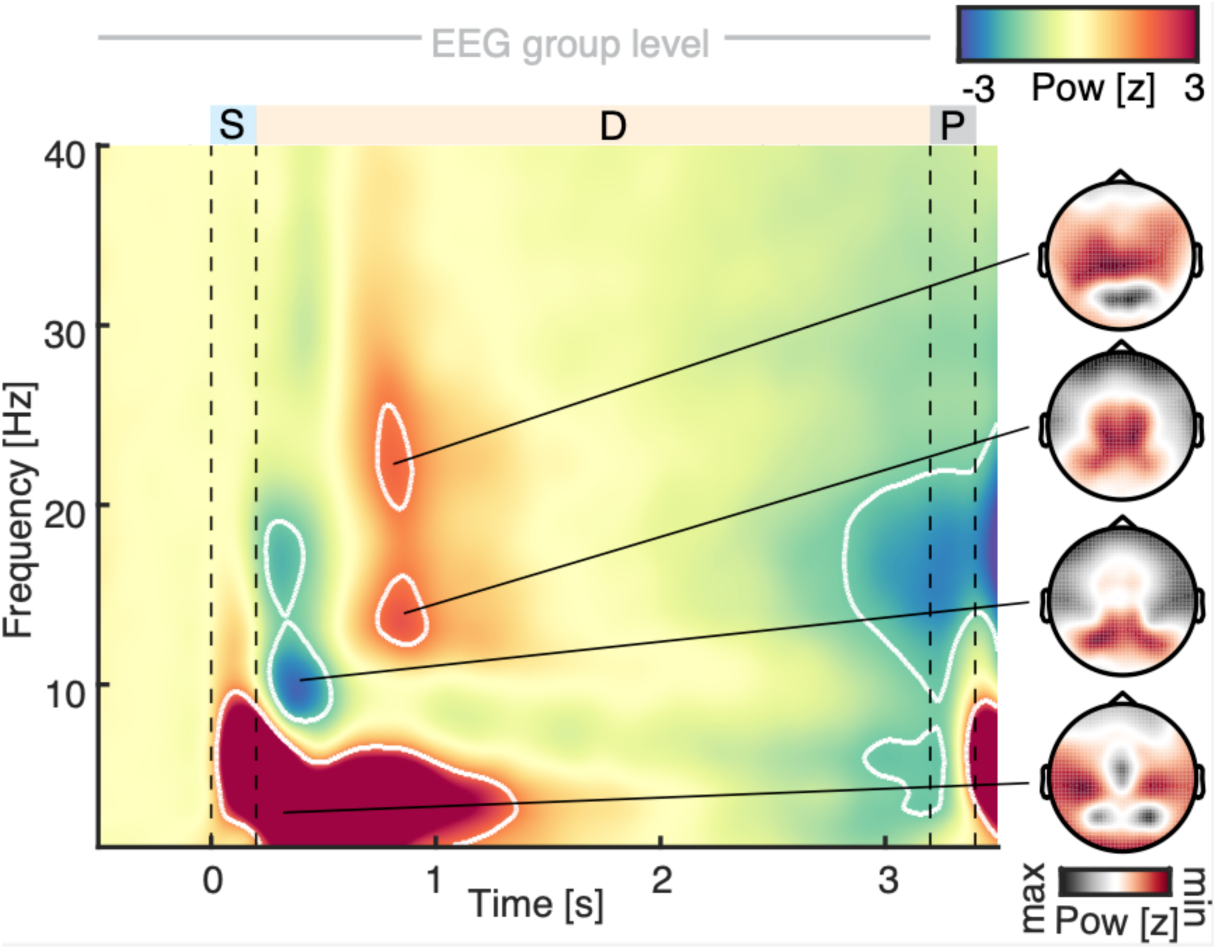
Grand average spectral power across all conditions and sessions. Left – temporal evolution. Cold colours - decrease, warm colours - increase compared to baseline (−0.5 to 0 s). White Contour marks a z-score of 1.65 (equivalent to p < 0.05, one-tailed). Note the low frequency power increase (0-7 Hz) starts from stimulus presentation, in line with an ERP effect. In the delay period (0.3 to 3.1 s), an alpha/beta (7-19 Hz) power decrease as well as a low beta (14-18 Hz) and high beta (18-25 Hz) power increase emerge. Right – spatial extent of power changes (black – maximal, red – minimal z-scored power). Note, the localization of alpha/beta effects to visual cortex and of beta effect to parietal and motor cortices.

**Figure 3:**
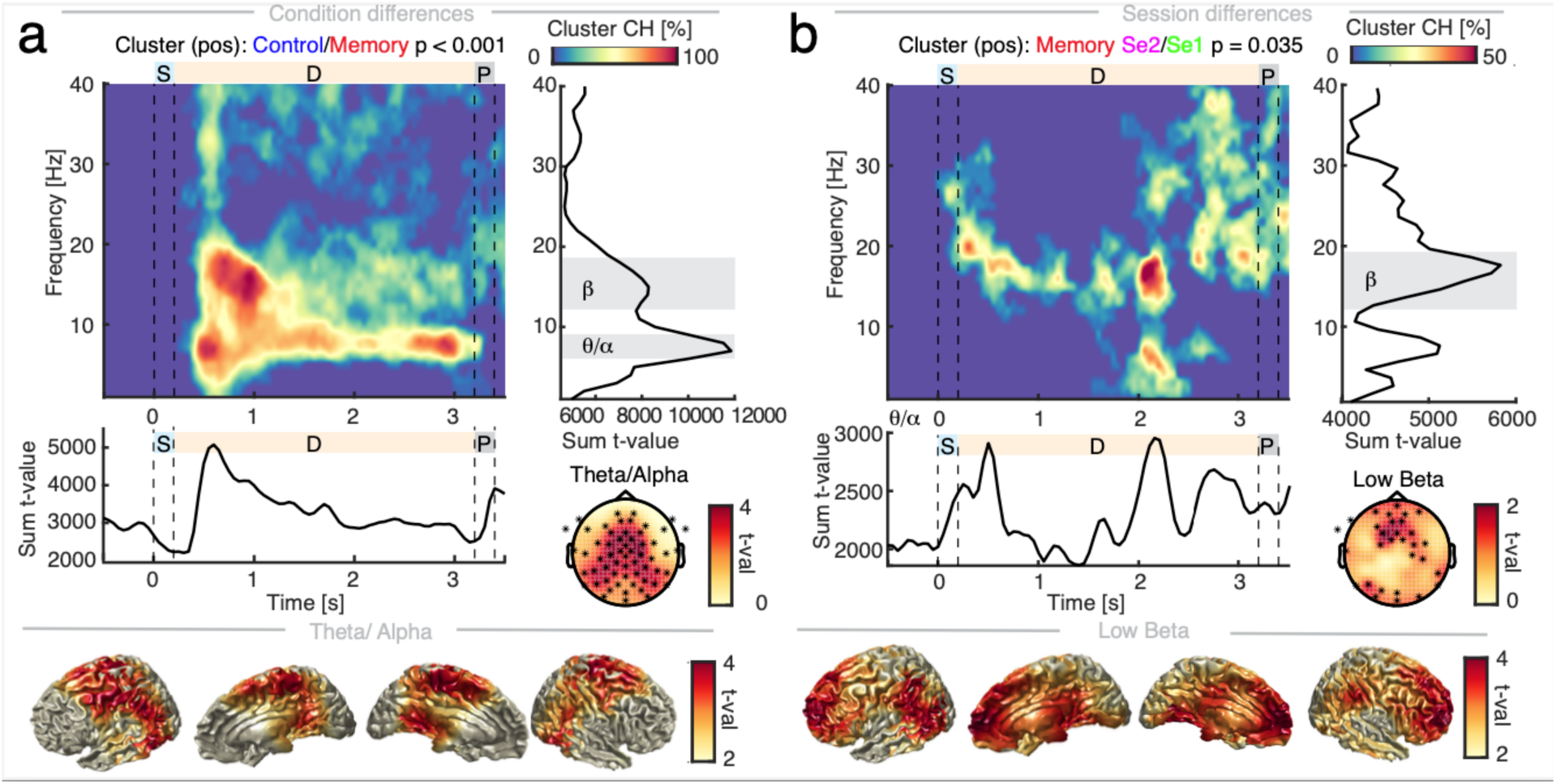
Spectral power changes across conditions and sessions. Sequence of WM task: blue/S – sample, orange/D – delay, grey/P – probe. **a**, Left upper – Condition power differences between control and memory averaged across sessions (baseline-corrected; n = 26). Colour: percent of channels in cluster from blue – 0 to red – 100%. Right upper – Sum of t-values per frequency. Note the peaks in theta/alpha (7 – 9 Hz) and low beta (14 –18 Hz). Left middle – Sum of t-values per time. Note the increased activity in theta/alpha in the control condition compared to the memory one, revealing a sustained suppression in memory trials. Right middle – spatial extent of theta/alpha power differences on scalp. Lower panel – source-interpolated on a template brain in MNI space. Low beta power exhibited a similar spatial spread and was therefore omitted here. Colour-coded: yellow – no difference to red – strong difference. **b**, Left upper – Memory session power differences (baseline-corrected; n = 26). Colour-coded: percent of channels in cluster from blue – 0 to red – 50%. Right upper – Sum of t-values per frequency. Note the peak in low beta (14 – 18 Hz). Left middle – Sum of t-values per time. Note the pulsing of beta during delay. Right middle – spatial extent of low beta power differences on scalp. Lower panel – source-interpolated on template brain in MNI space. Colour-coded: yellow – no difference to red – strong difference.

**Figure 4:**
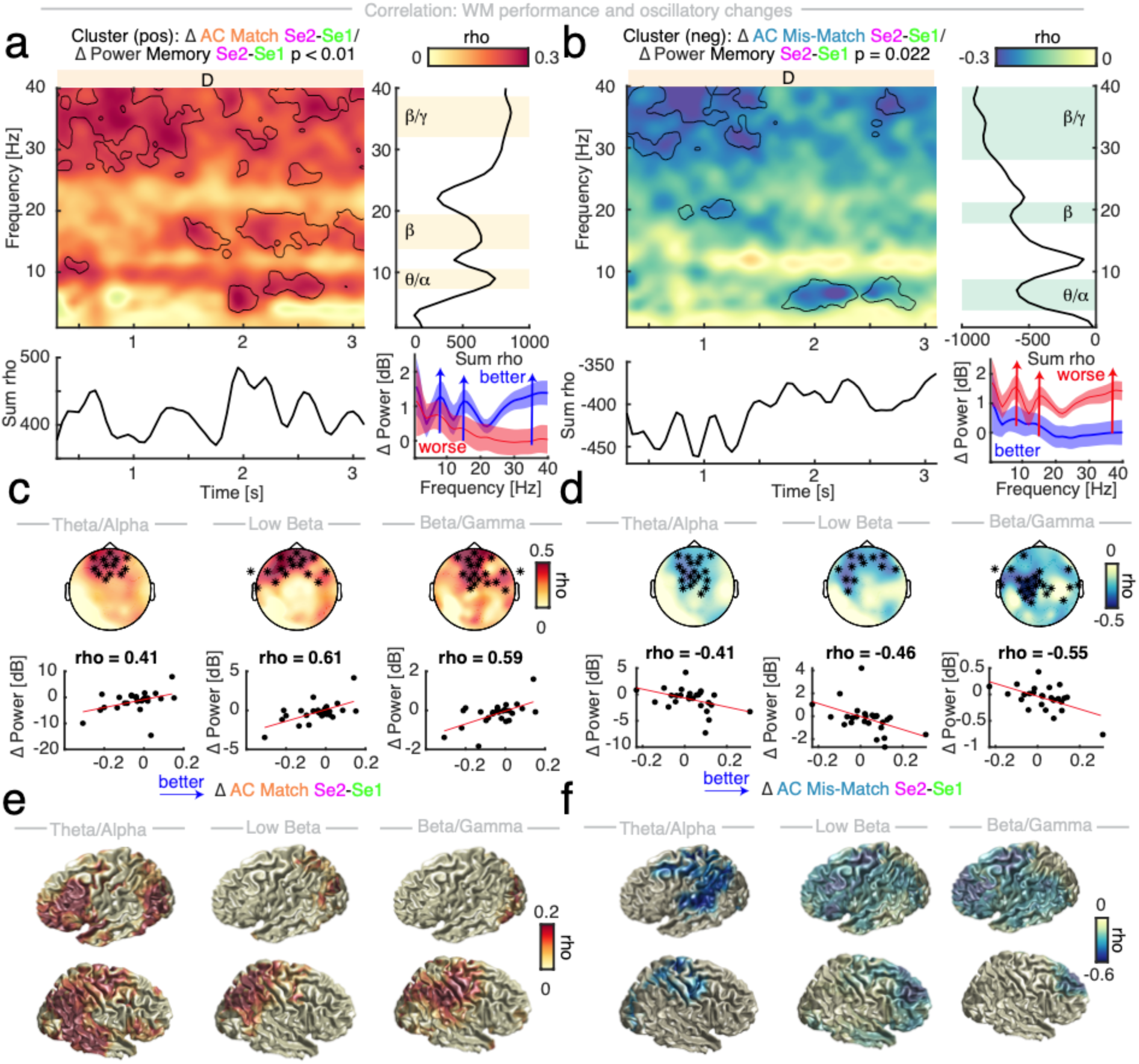
Behaviourally relevant oscillatory spectral changes. Association between memory session spectral power (Δ Se2 – Se1) and accuracy (AC) differences in the delay (0.3 to 3.1 s, n = 26). **a**, Match: Left upper – cluster permutation correlation. Black contour – rho > 0.2. Orange/D – delay. Right upper – Sum of t-values per frequency. Note the peaks in theta/alpha and low beta frequency range as well as a broadband effect with a peak in the high beta/low gamma range. Lower left – Sum of t-values over time. Lower right – Median AC Match split of power difference (Δ Se2 – Se1; baseline-corrected; mean with SEM). Red – worse, blue – better performance. **b**, Mis-Match: Left upper – cluster permutation correlation. Black contour – rho < -0.2. Orange/D – delay. Right upper – Sum of t-values per frequency. Note the negative peaks in similar frequency ranges. Lower left – Sum of t-values over time. Lower right – Median split AC Mis-match of power difference (Δ Se2 – Se1; baseline-corrected; mean with SEM). Red – worse, blue – better performance. Correlation of AC and power in the theta/alpha (7 – 9 Hz), low beta (14 – 18 Hz) and high beta/low gamma range (34 – 38 Hz). **c**, AC Match: Upper row – Spatial extent, Lower row – correlation rho. **d**, AC Mis-match: Upper row – Spatial extent, Lower row – correlation rho. **e**, AC Match: Rho-values source-interpolated on a standard template brain in MNI space (threshold p < 0.05). **f**, AC Mis-match: Rho-values in MNI space.

**Figure 5:**
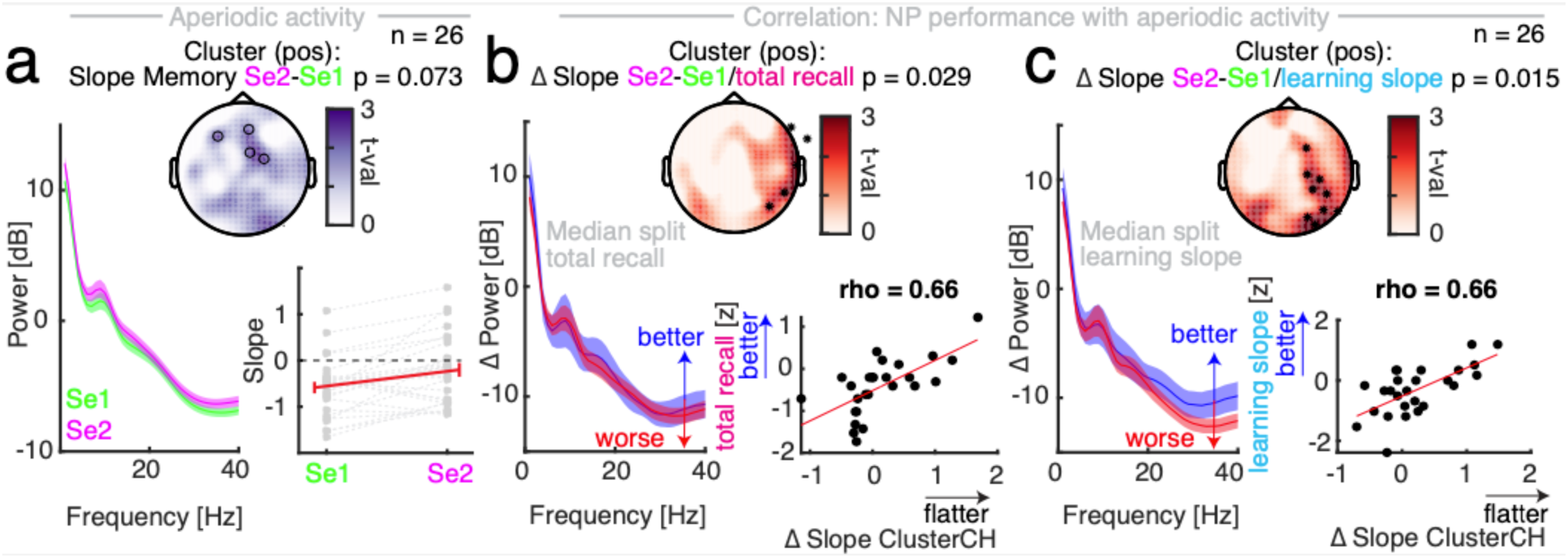
Aperiodic activity in pre- and post-anaesthesia memory sessions. **a**, Lower left – Spectral power. Middle – Spectral slope session difference borders on significance (*cluster permutation t-test* t_25_ = 9.6, p = 0.073) with a focus on frontal channels (black circles). Lower right – Spectral slopes (*post-hoc t-test:* t_25_ = 1.97, p = 0.060, d = 0.31). Grey dotted lines – single subjects. Red line – mean ± SEM. Association between session differences (Δ Se2 – Se1) of spectral slope and neuropsychological test scores (negative z-scores indicate worse performance; n = 26). **b**, Lower left – Median split of spectral power difference by total recall z-scores. Blue – better, red – worse performance. Middle - Spatial extent of and right lower – correlation between Δ slope with z-score total recall in cluster channels. **c**, Lower left – Median split of spectral power difference by learning slope z-scores. Blue – better, red – worse performance. Middle - Spatial extent of and right lower – correlation between Δ slope and z-score of learning slope in cluster channels.

Between conditions, we found that spectral power was significantly higher in the control conditions (*cluster-based permutation t-test:* t_sum_ = 6.53*10^4^, p < 0.001) in the theta/alpha (7 – 9 Hz) and low beta band (14 – 18 Hz), revealing a sustained suppression of theta/alpha power in the memory conditions (**Fig. 3a**, single sessions see **Figures S4**).

Between memory sessions (**Fig. 3b**), power significantly increased post-anaesthesia (*cluster-based permutation t-test:* t_sum_ = 2.16*10^4^, p = 0.035) in the low beta band (14 – 18 Hz). There were no differences between control sessions (*cluster-based permutation t-test:* p = 0.401). Importantly, this effect could not be explained by ERP differences (**Fig. S5**). The power increase in the low beta frequency band as well as in the theta/alpha and high beta/low gamma ranges were significantly correlated with a better accuracy in detecting match trials (**Fig. 4a**; *cluster-based permutation correlation:* t_sum_ = 5.89*10^4^, p < 0.001) but with a worse performance in recognizing mis-matches (**Fig. 4b**; *cluster-based permutation correlation:* t_sum_ = -3.85*10^4^, p = 0.022). Note, that power changes in these frequency bands were not correlated with the NP scores (**Fig. S3c,e**, *cluster-based permutation correlation with power session difference:* total recall p = 0.439, learning slope p = 0.358).

### Increased aperiodic activity is associated with better performance in neuropsychological assessment

In line with an increase of aperiodic activity, the PSD became flatter (i.e. the slope decreased) in the post-anaesthesia memory session (**Table 2, Fig. 5a**; *cluster-based permutation t-test:* t_sum_ = 9.60, p = 0.073).

A flattening of the PSD was associated with a better performance in the NP CVLT total recall (**Fig. 5b-c**; *cluster-based permutation correlation:* Δ slope/z-score total recall r = 0.66, p = 0.029) and learning slope (*cluster-based permutation correlation:* Δ slope/z-score learning slope r = 0.66, p = 0.015) but not with the WM performance (**Fig. S3f**; *post-hoc correlations*: Δ slope of cluster channels/ Δ AC match r = 0.19, p = 0.365; Δ slope of cluster channels/Δ AC mis-match r = 0.03, p = 0.876). Beta power increases were not associated with slope decreases (14 – 18 Hz; *post-hoc correlation*: r = 0.15, p = 0.449; **Fig. S3d**).

## Discussion

The current study examined the impact of surgery and general anaesthesia on postoperative cognitive performance and electrophysiology using neuropsychological (NP) assessment and a visual working memory (WM) task with concomitant scalp EEG. We observed a striking double dissociation between cognitive performance and EEG signatures. Behavioural results revealed that patients’ working memory performance in general did not change despite surgery under general anaesthesia.^27–29^ However, performance differences between match and mis-match trials emerged in the postoperative session. Performance increases in match trials in the WM task were associated with an enhancement of low and high beta oscillations in the postoperative session. Additionally, better cognitive function in NP assessment was accompanied by a rise of aperiodic, non-oscillatory brain activity. These findings reveal that rhythmic and arrhythmic neural activity tracks distinct facets of peri-anaesthesia cognition.

### Oscillatory signatures of working memory

Neural oscillations in various frequency bands have been implicated in WM.^13^ While theta (3-8 Hz) oscillations have been related to successful recognition^30^ and are thought to temporally structure WM items^13^, alpha oscillations (8-12 Hz) are often associated with inhibition^31, 32^ and assumed to suppress task-irrelevant activity.^13^ Beta activity (13-30 Hz) reflects at least two functionally and spatially distinct components, the sensorimotor beta oscillation that mediates motor control and frontal beta activity, which is related to top-down cognitive processing.^27, 28^ Increased beta and gamma activity (∼40 Hz)^10^ have been linked to the active maintenance of information in WM.^13, 20, 28^ Here, we found a memory-related suppression of theta/alpha and low beta activity (7-18 Hz; **Fig. 3a**) over fronto-parietal cortices in line with a release from inhibition in task-relevant areas.^31, 33^

Between pre- and post-anaesthesia memory sessions, low beta (14 – 18 Hz) oscillations increased, occurring in bursts over frontal cortex during the entire delay period (**Fig. 3b**) potentially signalling top-down control.^27–29^ Importantly, this beta enhancement was associated with differential processing of match and non-match trials in the WM task (**Fig. 4**). Increases in performance in match trials from pre- to postoperative assessments were associated with increases in theta (7-9 Hz), low beta (14-18 Hz), and beta/gamma (34-38 Hz) oscillatory activity. Whereas decreases in performance in non-match trials from pre- to postoperative assessments were associated with increases in theta (7-9 Hz) and beta/gamma (34-38 Hz) oscillatory activity (**Fig.4**). Beta and gamma bursts during WM maintenance were suggested to be related to readout and control mechanism in WM tasks^34^ and also with inhibition of competing visual memories.^35^ Thus, our findings are well in line with theories proposing that enhanced oscillatory synchrony enables the efficient network communication which is the fundament of optimal cognitive processing.^36^

### The importance of aperiodic activity for cognitive function

While some recent evidence suggests that a shift of oscillatory peak frequency of the posterior alpha frequency activity can outlast anaesthesia and might relate to cognitive performance^37^, the importance of aperiodic activity for cognitive function in a perioperative setting has not yet been investigated. Recently, several lines of inquiry highlighted that aperiodic activity tracks task-related neural activity in a variety of cognitive domains such as attention^18^, motor execution^38^ and working memory^14, 39^ and is related to better task performance. In the present study, we observed a similar pattern, namely that enhanced aperiodic activity (i.e. flattening of the PSD/ decrease of spectral slope) after anaesthesia was associated with a better performance in the neuropsychological assessment (**Fig. 5**). If aperiodic activity was not enhanced or even reduced after anaesthesia, patients performed worse (**Fig. 5**). Our findings demonstrate that an increase of aperiodic activity tracks a selective activation of task-relevant cortices resulting in better cognitive performance.

### Anaesthetic-induced inhibition overhang as a potential contributor to postoperative neurocognitive impairment

Experimental evidence and computational modelling have linked aperiodic activity to the excitation to inhibition balance of the underlying neuronal population, where an increase of inhibition (e.g. by GABAergic drugs like propofol) results in a steeping of the PSD (i.e. an increase of spectral slope).^15–17^ Increased cortical (excitatory) activity, on the other hand, led to a flattening of the PSD (i.e. a decrease in slope) that was associated with improved task performance.^18, 19^ Computational modelling established that aperiodic activity indexes the underlying balance between excitatory and inhibitory neuronal activity on the population level^17^. Anaesthetics like propofol and sevoflurane increase cortical inhibition via GABA receptors. Recent electrophysiological studies confirmed that increased inhibition reduces aperiodic activity as indexed by a steepening of the PSD, i.e. an increase of spectral slope.^15–18^

To date it remains unclear how fast this inhibition dissipates after drug administration is discontinued. While the healthy young brain is resilient to electrophysiological disturbances by anaesthesia^40^, elderly patients are more sensitive to anaesthetics and therefore more likely to enter burst suppression^4^ - a pattern of brain activity induced by a maximum of inhibition where bursts of activity are followed by periods of neuronal silence.^41^ Time spent in that (too) deep plane of anaesthesia is an independent risk factor for developing POD.^2, 42^ Not only age but also individual brain vulnerability plays a role in susceptibility to adverse cognitive outcomes after anaesthesia: Patients that reacted with intraoperative EEG suppression at lower doses of anaesthetics were more likely to develop POD.^43, 44^ In addition, a recent study that examined EEG parameters before, during and after anaesthesia found that a lower preoperative spectral edge frequency and gamma power were independently related to the development of POD.^5^ Note that these findings could also be explained by a steeper slope of the preoperative PSD (i.e. more inhibition) in the POD group compared to the patients that did not develop POD. In the current study, patients that showed an increase of aperiodic activity (less inhibition) after anaesthesia, performed better in postoperative neuropsychological assessment whereas patients that exhibited a decrease (more inhibition) performed worse (**Fig. 5**).

Taken together, these findings suggest that anaesthesia-induced inhibition potentially outlasts drug administration in elderly patients and that this inhibition overhang impedes optimal postoperative cognition. Pre-existing alterations of neuronal balance between excitation and inhibition (such as a premedication with e.g. benzodiazepines) could be a predisposing factor for developing postoperative cognitive decline. Thus, aperiodic brain activity might provide a valuable marker to track perioperative cognition - before, during and after anaesthesia.

### Strengths and limitations

Key advantages of the current study are: 1) It focused on electrophysiological changes in the early postoperative period. 2) Both NP assessments as well as a WM task were obtained. 3) It had a substantial scalp EEG coverage (62 channels instead of 10 or 20 sensors). 4) The WM task featured a control condition to evaluate memory-specific changes. 5) It examined an elderly cohort vulnerable to perioperative cognitive changes. The study had the following limitations: 1) It was conducted at a prostate cancer centre; thus, only men were included limiting generalisability of results. 2) There was no matched control group, restricting our ability to differentiate session-from purely anaesthesia-related changes. 3) Fifteen patients did not complete the second session, potentially patients that suffered from more postoperative complications. High drop-out rates are frequently reported in studies focusing on perioperative cognitive disorders (Ref) and pose a relevant source of selection bias. To address this issue, we compared demographic and clinical characteristics between patients, who completed all pre- and postoperative assessments and those who did not without observing relevant differences.

## Conclusion

Here, we present empirical evidence that oscillatory and aperiodic dynamics track different aspects of perioperative cognition. Specifically, aperiodic activity as an index of population activity and anaesthesia-induced inhibition overhang might prove to be a valuable biomarker in tracking cognitive function in the perioperative setting.

## Acknowledgments

The authors would like to thank Randolph Helfrich, Hertie-Institute for Clinical Brain Research, Tübingen, for his advice regarding data analysis as well as his valuable input on the manuscript.

## Details of author’s contribution

All authors declare that they took part in the revision of this manuscript, that they approved of the final version and they agree to be accountable for all aspects of the work. Furthermore, the authors contributed in the following areas: JDL: Data analysis, data interpretation, writing of first draft of manuscript; UH: Patient recruitment, data collection; JD, AKE, CZ: Study design; TS: Study design, data collection, data analysis and interpretation; MF: Study design, patient recruitment, data collection, data analysis and interpretation.

## Data and custom code availability

Data will be made available upon reasonable request to Marlene Fischer. Custom code will be shared upon reasonable request to the corresponding author.

## Supplemental Information

### Supplemental Material and Methods

#### Flow Diagram of Study

**Fig. S1:**
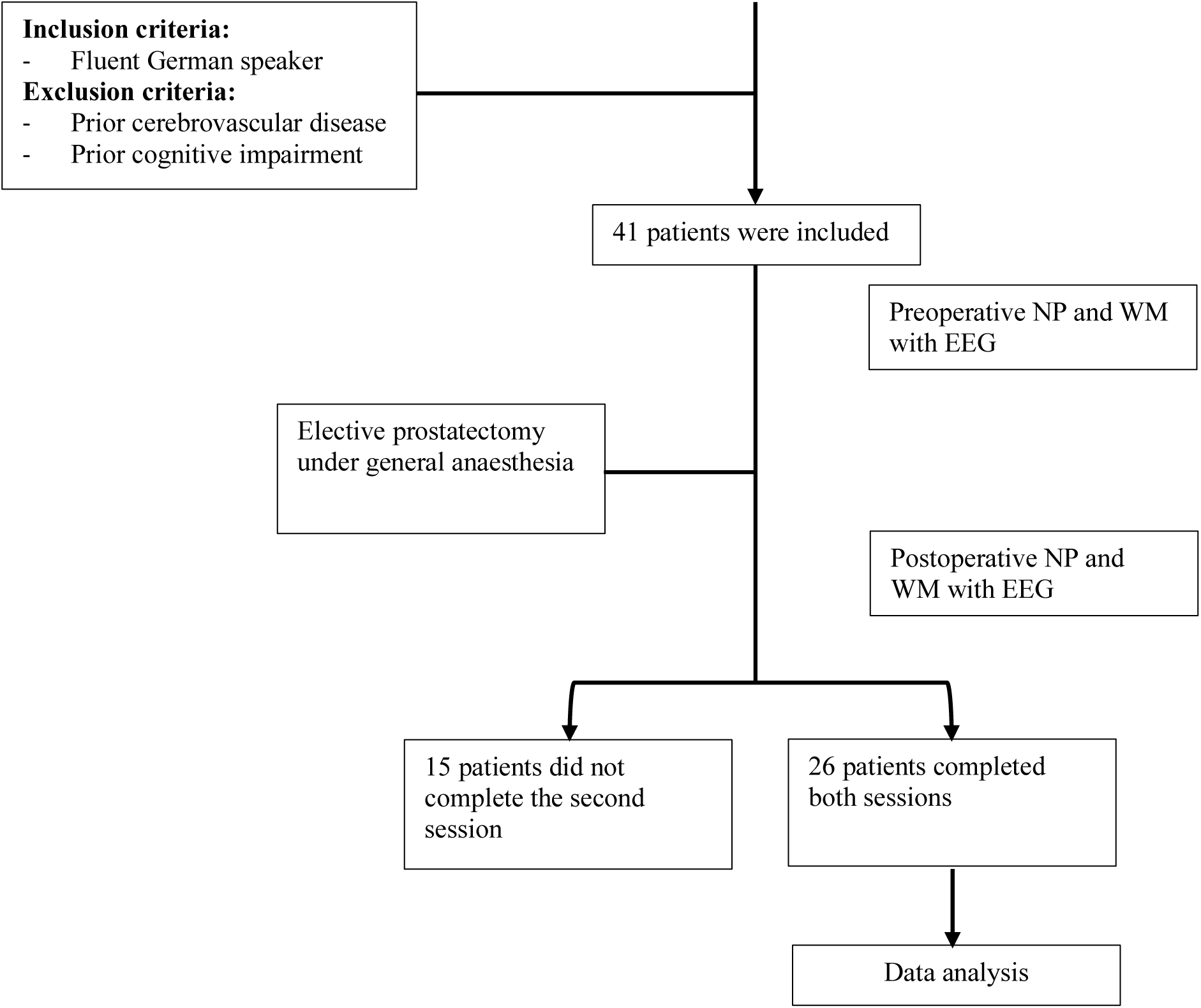
Flow Chart.

#### Working Memory Task

Each trial of both memory and control condition began with a fixation dot, followed by a 200 ms sample presentation and a 3000 ms delay period in which the fixation dot reappeared. In the memory condition, patients were then presented with the probe for 200 ms (**Fig. 1**) and had to indicate whether the probe matched the sample (match) or not (non-match). Responses were registered by pressing one of two buttons using either their index or middle finger (counterbalanced). The inter-trial interval was randomly jittered in steps of 100 ms ranging from 2.5 to 3 seconds. In the control condition, an ellipse was presented for 50 ms instead of a second image. Here, patients had to report whether the ellipse was vertically or horizontally oriented.^15^ Performance in the control condition was matched to the memory condition by changing the shape and luminance of the ellipse.^15^ In addition, in 20% of control trials, the contrast and shape were chosen randomly to disguise the relationship to memory performance.^15^ Before the beginning of the recording, patients had a training session where 4 trials of each condition were presented that were not part of the consecutive task.^15^

#### Anaesthesiologic management

General anaesthesia was induced with sufentanil (0.3–0.7 μg/kg) and propofol (2–3 mg/kg) followed by neuromuscular blockade with rocuronium (0.6 mg/kg) to facilitate endotracheal intubation. A gastric tube was inserted in all patients and prophylactic antiemetic medication was administered preoperatively (dexamethasone 4 mg). During surgery, rocuronium was titrated under the guidance of neuromuscular blockade monitoring (train-of-four, TOF-Watch Organon; IntelliVue NMT module, Philips GmbH, Hamburg, Germany). Anaesthesia depth was monitored with a bispectral index monitor (BIS™, Medtronic GmbH, Meerbusch, Germany). Sevoflurane-sufentanil was used for anaesthesia maintenance to achieve an end-tidal sevoflurane concentration of 2.0 vol% (MAC 0.8–1.2). To maintain normothermia we used a forced-air warming system throughout the entire procedure. Patients who underwent robot-assisted surgery received peritoneal insufflation with carbon dioxide and were positioned at a 45-degree head-down tilt. Postoperative pain management included non-opioid medication (metamizole 1000 mg/100 ml) 30 minutes before emergence and every 4–6 hours thereafter. In the post-anaesthesia care unit (PACU) piritramide 3.75–7.5 mg was administered intravenously when pain scores exceeded 3. Subsequent PACU management in all patients included frequent control of wound drains and postoperative urine output, blood gas analyses, and pain management as described above.

#### Data cleaning – Excluded channels and trials

In total, 56 channels of Session 1 (not including EOG leads; 3.47 %) and 68 channels for Session 2 (4.29 %) were excluded, averaging to 2.15 channels/subject in Session 1 and 2.62 channels/subject in Session 2. Excluded channels were interpolated using their neighbours (ft_channelrepair). Regarding trials, a total of 194 trials of Session 1 (3.59 %) and 172 trials of Session 2 (3.18 %) were identified as noisy and removed, averaging to 7.46 trials/subject for Session 1 and 6.62 trials/subject for Session 2.

#### Event-Related Potential (ERP) analysis

For ERP analysis, electrophysiological data of each session was low-pass filtered below 30 Hz and split into memory and control conditions. Data was then time-locked to the respective baseline (−200 - 0 ms before sample presentation) and averaged across trials. For a comparison between conditions, we averaged across sessions (**Fig. S5**).

#### Beamformer analysis

We source-localized power differences in frequency bands of interest and their significant correlation with WM performance (Fig. 3, 4). Cortical sources of the sensor-level EEG data were reconstructed by using a LCMV (linearly constraint minimum variance) beamforming approach ^20^ to estimate the time series for every voxel on the grid. A standard T1 MRI template and a BEM (boundary element method) headmodel from the FieldTrip toolbox^19^ were used to construct a 3D template grid at 1cm spacing in standard MNI space. Electrode location were positioned on a standard head and aligned with MNI space using the FieldTrip toolbox^19^. Prior to source projection, sensor level data was common average referenced and epoched into 5 second segments. To minimize computational load, we selected a two second data segment (1 to 3 seconds) from the delay period of every epoch to construct the covariance matrix. The LCMV spatial filter was then calculated using the covariance matrix of the sensor-level EEG data with 5% regularization. Spatial filters were constructed for each of the grid positions separately to maximally suppress activity from all other sources. The resulting time courses in source space then underwent spectral analysis using a Fourier transform using a multitaper approach (‘*mtmfft*’ of ft_freqanalysis from FieldTrip using ‘dpss’ taper, a smoothing frequency +/- 1 Hz and a 0,5 second moving time window). For the memory/control comparison, power spectra were averaged across sessions prior to statistical analysis. Power differences between both memory and control and well as between memory sessions, and their correlation with behaviour were then tested using cluster-based permutation approaches employing either repeated t-tests or correlation (see below for details). The results were calculated at every voxel in source space, converted to z-values and then interpolated onto a standard template brain in MNI space.

### Supplemental Results

#### Patient characteristics

**Table S1.**
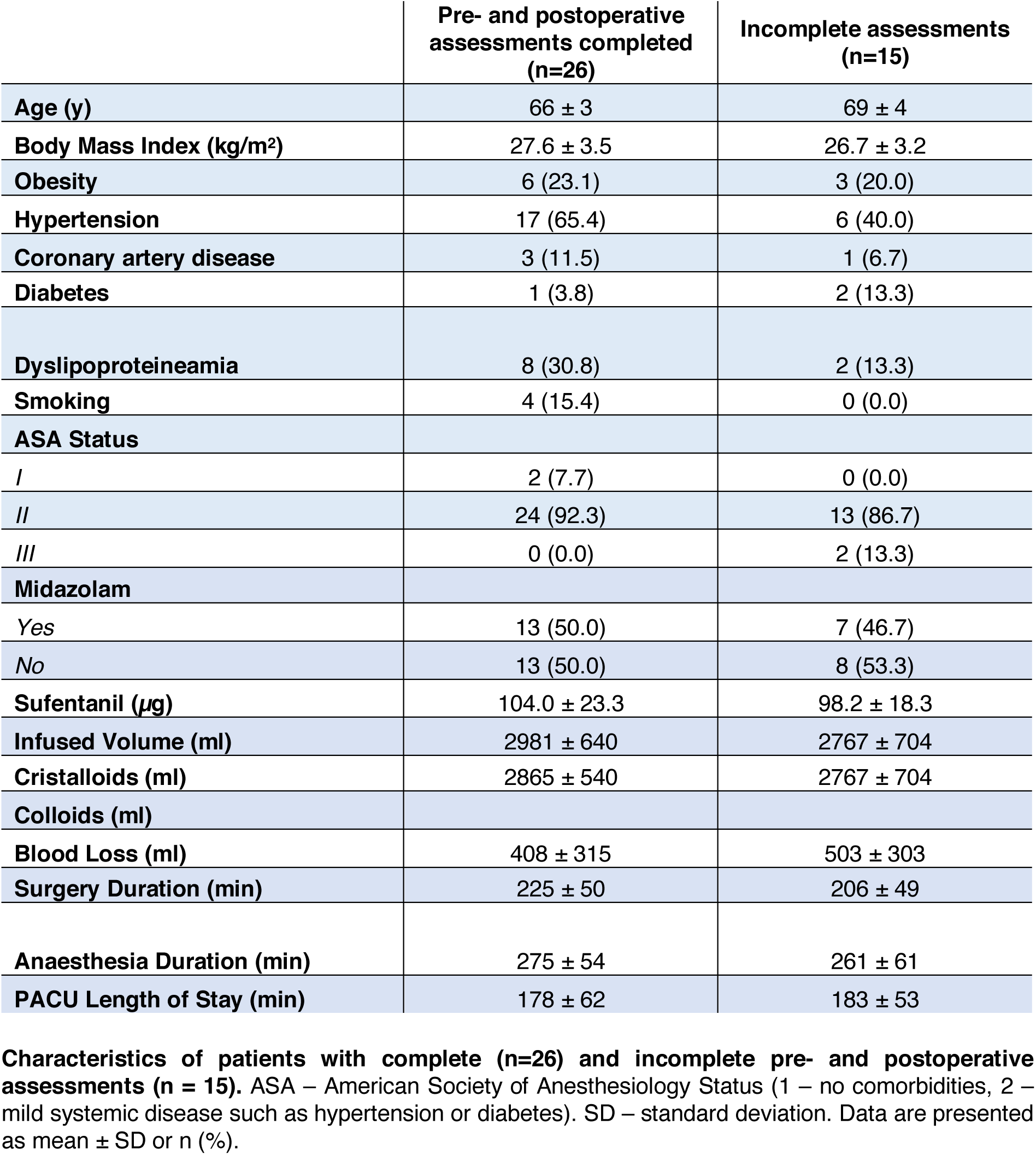

#### Behavioural Performance – Neuropsychological Assessment

**Table S2.**

|  | Se 1 |  | Se 2 |  | z-score |  |
| --- | --- | --- | --- | --- | --- | --- |
|  | Mean | SD | Mean | SD | Mean | SD |
| **Total recall** | 44 | 9 | 40 | 9 | -0.42 | 0.66 |
| **Learning slope** | 1 | 0 | 1 | 0 | -0.31 | 0.81 |
| **TMT_B (s)** | 80 | 22 | 73 | 20 | -0.17 | 0.54 |
| **GPT_dom (s)** | 82 | 8 | 80 | 11 | -0.08 | .055 |
| **Digit Span** | 5 | 1 | 5 | 1 | -0.04 | 0.79 |

**Absolute values and z-scores of neuropsychological assessments (n = 26).** Se – session. TMT – Trail Making Test. GPT – Grooved-Pegboard Test. dom – dominant. Data are presented as mean ± SD.

#### Behavioural Performance – Working Memory Reaction times

In the WM task, there was no significant influence of condition or session on reaction times (RT; *repeated-measures ANOVA:* condition F_1,25_ = 0.366, p = 0.551; session F_1,25_ = 0.008, p = 0.928; condition*session F_1,25_ = 0.216, p = 0.646; **Table 2**). However, patients exhibited a tendency to be faster for the non-match than the match trials (see **Table S2**; *repeated-measures ANOVA:* match F_1,25_ = 3.785, p = 0.063; session F_1,25_ = 0.141, p = 0.710; match*session F_1,25_ = 0.929, p = 0.344).

#### Event-related potentials differentiate memory and control conditions

We could identify a total two clusters that marked significant differences on the group level: One late (*cluster-based permutation t-tests:* 1 to 3.1 s, t_1_ = -8.87*10^4^, p_1_ = 0.002, 37 channels with a focus on posterior parietal cortex) and one early in the delay period (reflecting the p300; 0.2 to 1 s, t_2_ = -4.74*10^4^, p_2_ = 0.004, 34 CH posterior parietal) mirrored by two positive ones (early 0.2 to 1 s, p_1_ = 0.014, t_1_ = 4.06*10^4^, 43 channels over frontal cortex; late 2.1 to 2.95 s, p_2_ = 0.016, t_2_ = 3.27*10^4^, 22 frontal channels; see **Fig. S5**).

When contrasting the ERPs of the post- and pre-anaesthesia memory sessions (Se2-Se1), there were no significant differences (*cluster-based permutation t-tests:* one positive cluster of 11 central channels, 0.57 to 0.9 s, p = 0.272, t = 5.14*10^4^). Note, that both the comparison of ERPs of control sessions or the memory-control differences did not show significant differences as well (Ctrl: positive cluster. 10 central channels, p = 0.296, t = 4.68 * 10^3^; Difference: positive cluster, 15 central channels, p = 0.805, t = 2.31 * 10^3^; **Fig. S5**).

## Supplemental Figures

**Figure S2:**
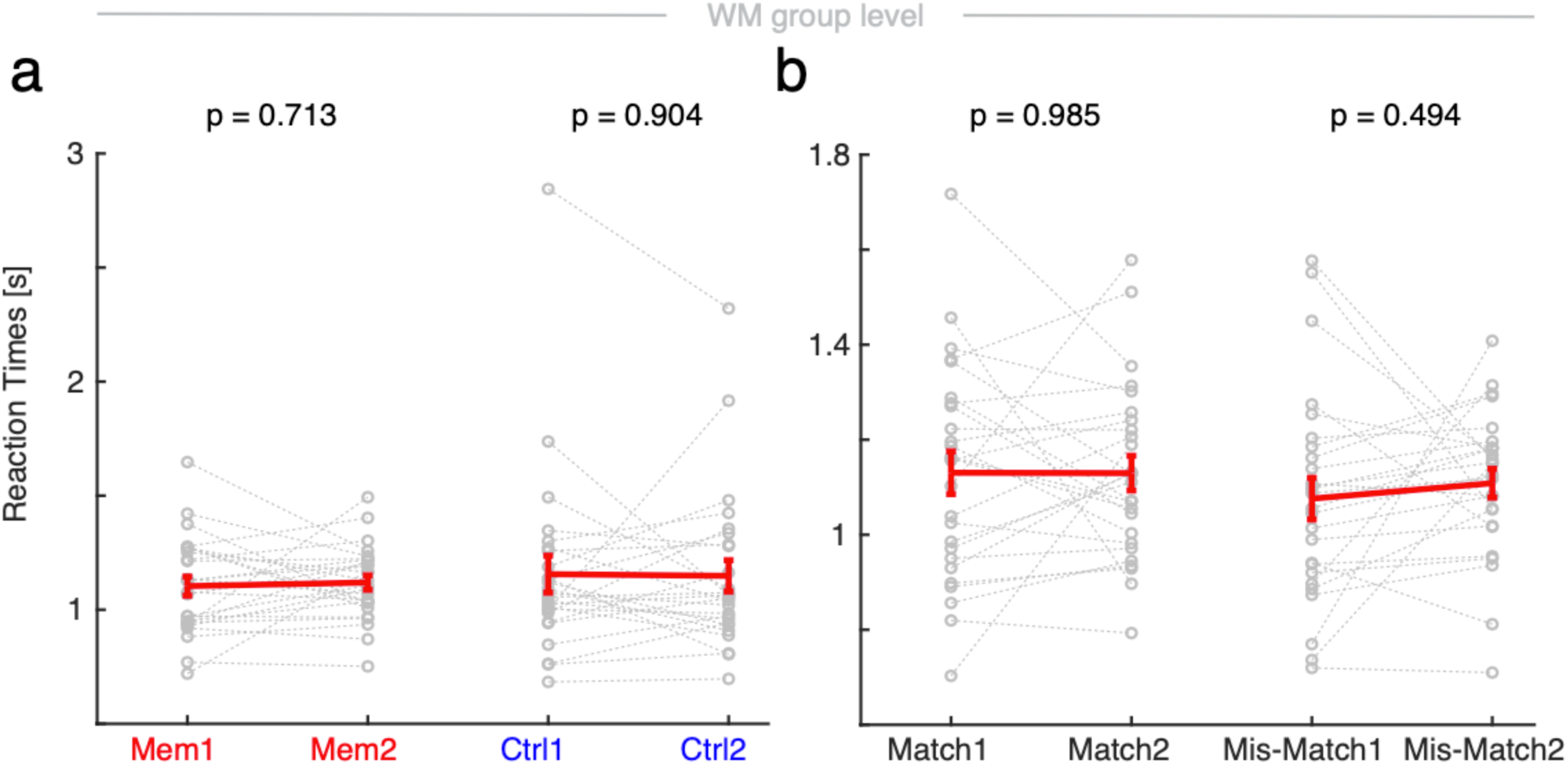
Behaviour in the visual working memory task. **a**, Reaction times in seconds (s) in memory and control condition before (1) and after (2) anaesthesia (n = 26). *Repeated-measures ANOVA:* condition F_1,25_ = 0.366, p = 0.551; session F_1,25_ = 0.008, p = 0.928; condition*session F_1,25_ = 0.216, p = 0.646. Grey dots – single subjects. Red line – mean ± SEM. P-values *post-hoc uncorrected t-tests*. **b**, Reaction times in seconds (s) between memory sessions in match and non-match trials. *Repeated-measures ANOVA:* match F_1,25_ = 3.785, p = 0.063; session F_1,25_ = 0.141, p = 0.710; match*session F_1,25_ = 0.929, p = 0.344. Grey dots – single subjects. Red line – mean ± SEM. P-values *post-hoc uncorrected t-tests*.

**Figure S3:**
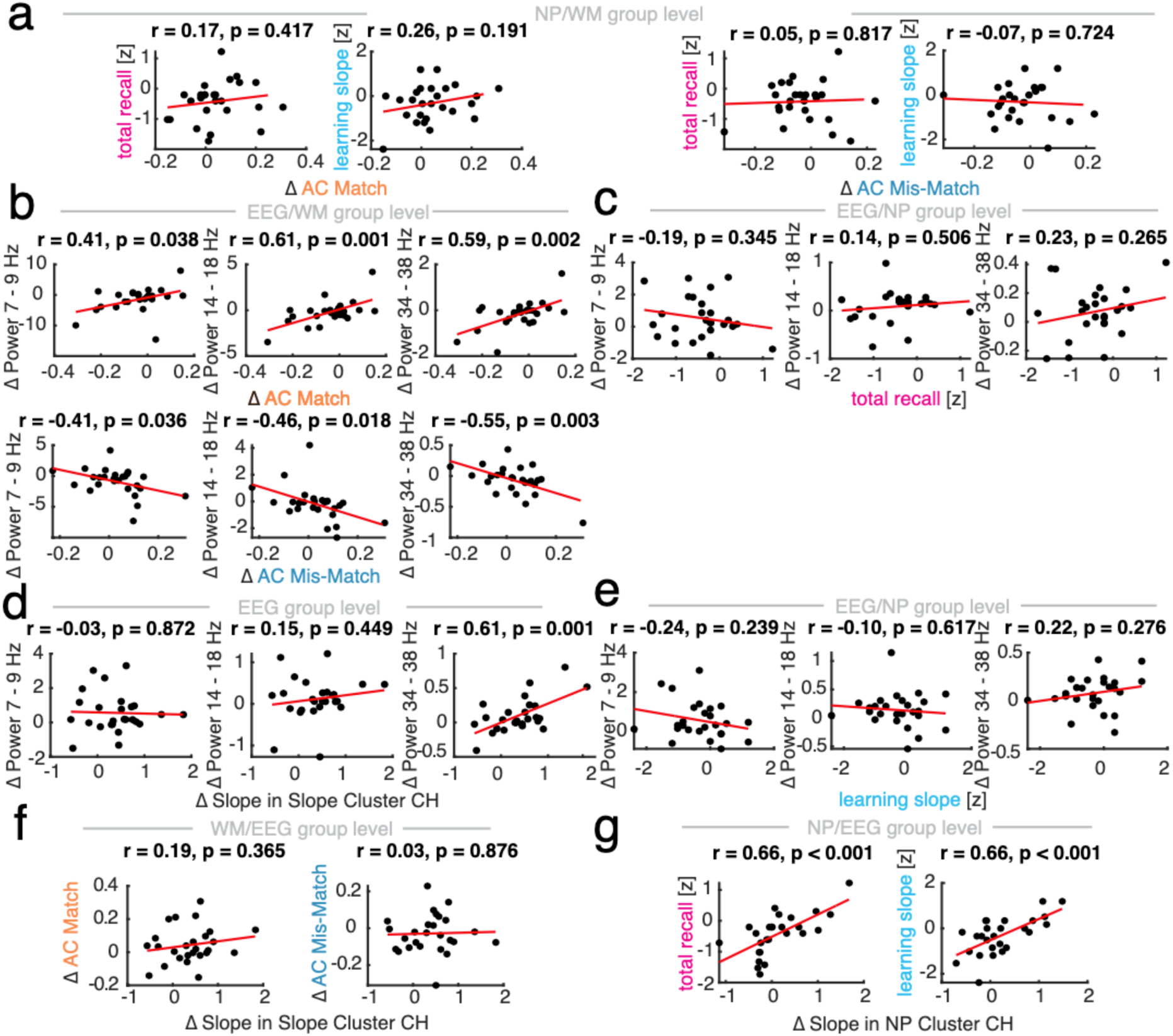
Associations between neuropsychological assessment, working memory performance, oscillatory and aperiodic electrophysiological features. **a**, Association of neuropsychological (NP) assessment (z-scores of California Verbal Learning Task (CVLT) total recall and learning slope) and session differences in performance between the memory sessions of the working memory (WM) task (Δ Se2-Se1, n = 26). Left – total recall and learning slope with Δ match accuracy; Right – Δ mis-match accuracy. **b**, Association of electroencephalogram (EEG) oscillatory power differences between memory sessions in frequency bands of interest (Δ Se2-Se1, n = 26) and WM Δ match and Δ mis-match accuracy in cluster channels (*post-hoc correlations*; session differences in spectral power and accuracy *cluster-permutation test:* Δ AC match p < 0.01, Δ AC mis-match p = 0.022; **Fig. 4**). **c**, Association of EEG oscillatory power differences between memory sessions in frequency bands of interest (Δ Se2-Se1, n = 26) and NP z-score of CVLT total recall. **d**, Association of EEG oscillatory power differences between memory sessions in frequency bands of interest (Δ Se2-Se1, n = 26) and EEG spectral slope (30 – 40 Hz) difference between memory sessions in cluster channels (Δ Se2-Se1 spectral slope *cluster-permutation test:* p = 0.073, **Fig. 5**). **e**, Association of EEG oscillatory power differences between memory sessions in frequency bands of interest (Δ Se2-Se1, n = 26) and NP z-score of CVLT learning slope (**Fig. 5**). **f**, Association between WM performance and EEG Δ spectral slope (Δ Se2-Se1, n = 26). Left – accuracy in match and Right – accuracy in mis-match discrimination in slope difference cluster channels. **g**, Association between NP performance and EEG Δ spectral slope (Δ Se2-Se1, n = 26). Left – z-score CVLT total recall and Right – z-score CVLT learning slope in NP cluster channels (Δ spectral slope and total recall *cluster-permutation test:* p = 0.029; Δ spectral slope and learning slope *cluster-permutation test:* p = 0.015; **Fig. 5**).

**Figure S4:**
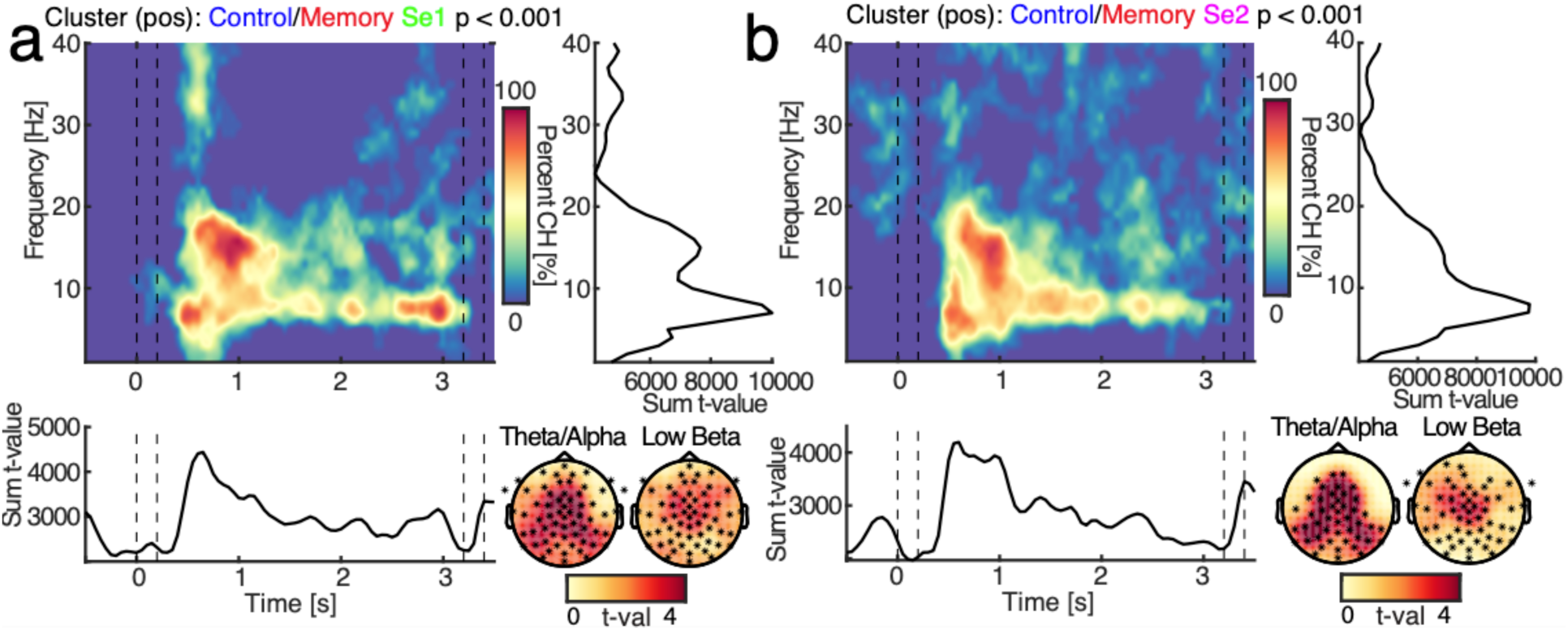
Spectral power changes in pre- and post-anaesthesia session. Sequence of WM task: blue/S – sample presentation, orange/D – delay, grey/P – probe presentation. **a**, Control and memory condition power differences (baseline-corrected) in the pre-anaesthesia session (Se1). Left upper panel – Spectral power changes between control and memory in session 1 (p < 0.001, n = 26). Colour-coded is the number of channels that are part of the cluster from blue – no channels to red – 100% of channels. Right upper panel – Sum of t-values per frequency. Note the peaks in the theta/alpha (7 – 9 Hz) and low beta (14 –18 Hz) frequency range. Left middle panel – Sum of t-values per time. Note the increased activity in the theta/alpha band in the control condition, revealing a sustained suppression in the memory session. Right middle panel – spatial extent of spectral power difference between conditions in the theta/alpha and low beta band. **b**, Post-(Se2) anaesthesia control and memory condition power differences (baseline-corrected). Left upper panel – Spectral power changes between control and memory in session 2 (p < 0.001, n = 26). Colour-coded is the number of channels that are part of the cluster from blue – no channels to red – 100% of channels. Right upper panel – Sum of t-values per frequency. Note the peaks in the that/alpha (7-9 Hz) and low beta (14 – 18 Hz) frequency range. Left middle panel – Sum of t-values per time. Right middle panel – spatial extent of condition difference in the theta/alpha (7-9 Hz) and low beta frequency (14 – 18 Hz).

**Figure S5:**
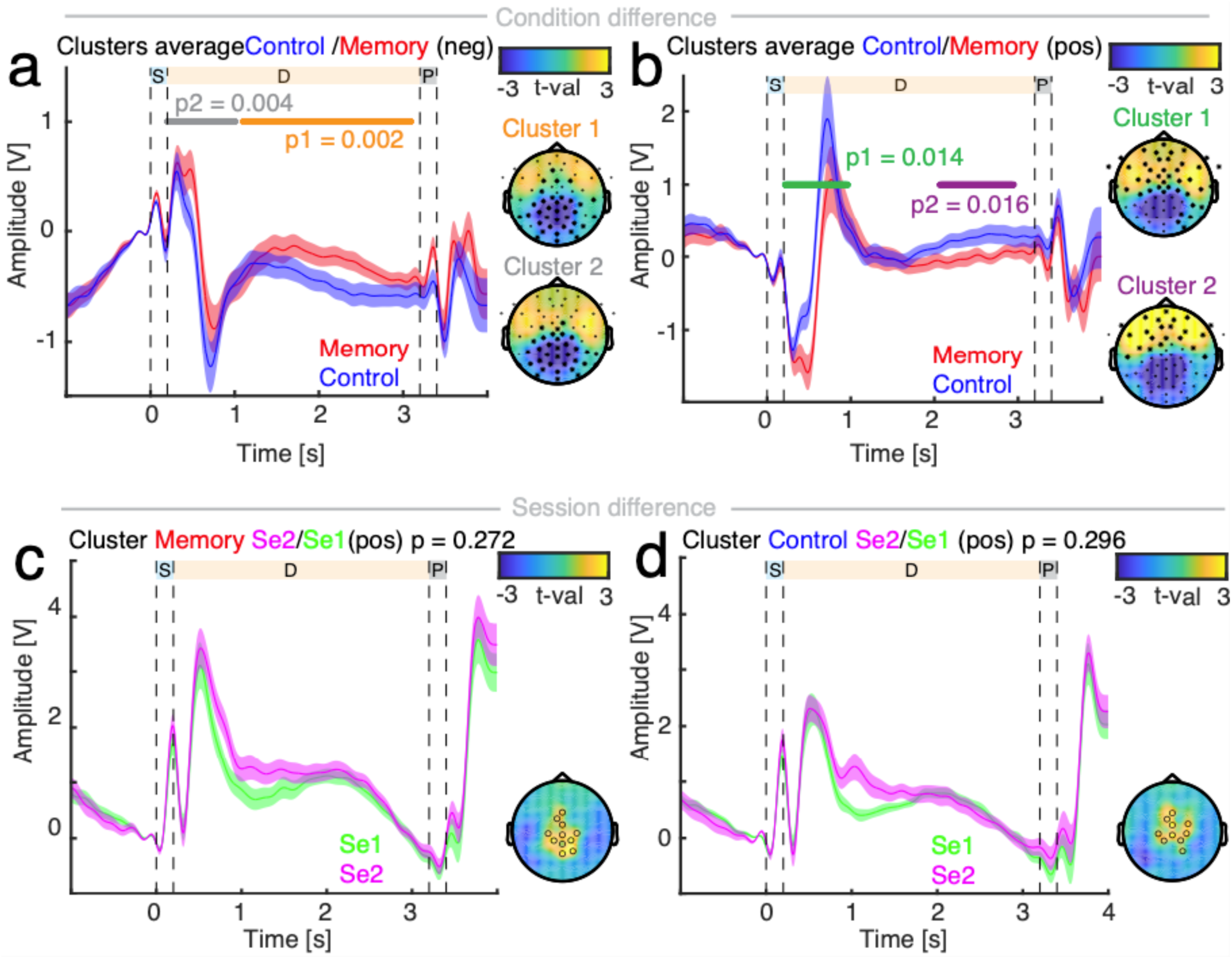
Comparison of condition and session Event-related potentials. Sequence of WM task: blue/S – sample, orange/D – delay, grey/P – probe. **a**, Temporal (left) and spatial (right) extent of condition differences of event-related potentials (ERP) between average control (blue) and memory (red; n = 26). Two negative, parieto-central clusters were significant: First one in delay (orange, p = 0.002), second one early (grey, p = 0.004) reflecting the ERP. **b**, Temporal (left) and spatial (right) extent of ERP differences between average memory (red) and control (blue). Two positive frontal clusters emerged: First one early (green, p = 0.014) reflecting the ERP, second late in delay (purple, p = 0.016). Note the mirrored effect compared to a. **c**, Pre- (green) and post-anaesthesia (magenta) memory sessions did not show significant temporal (left) or spatial (right) ERP differences. **d**, Pre- (green) and post-anaesthesia (magenta) control sessions did not show significant temporal (left) or spatial (right) ERP differences.

**Figure S6:**
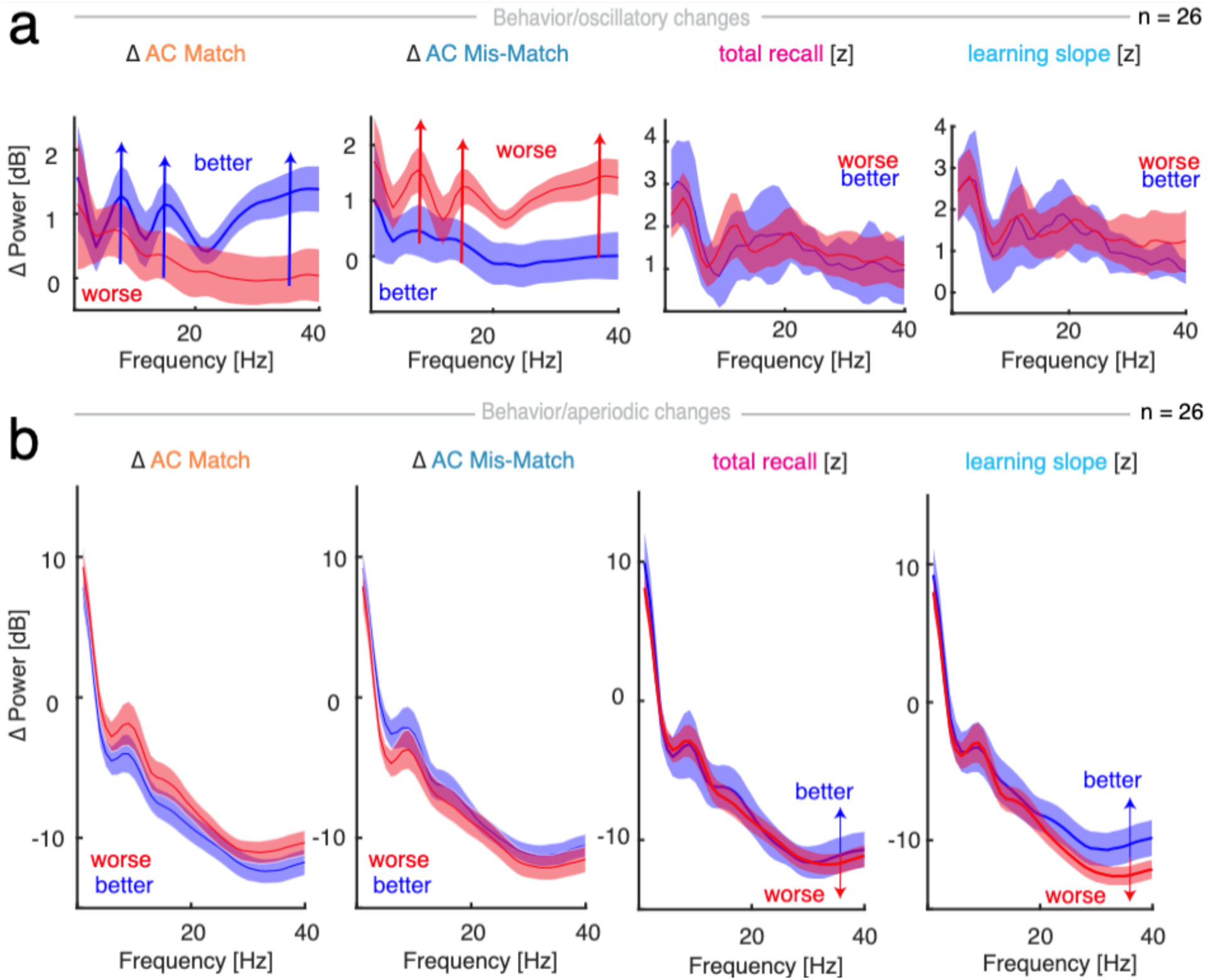
Electrophysiological signatures by median split of cognitive performances. **a**, Oscillatory changes: median split of baseline-corrected spectral power differences between post- and pre-anaesthesia sessions (Δ Se2 - Se1, n = 26). Left – split by median difference (Δ Se1-Se2) of Δ accuracy of match discrimination; left middle - Δ mis-match discrimination. Note, the power differences in certain frequency bands (theta, low and high beta). Right middle – Power split by median z-score of neuropsychological assessment by the California Verbal Learning Task (CVLT) total recall and right – learning slope. Blue – better performance than median, red – worse performance than median. Note, that there are no relevant oscillatory changes that track NP performance. **b**, Aperiodic changes: median split of spectral power differences between post- and pre-anaesthesia memory sessions (Δ Se2 - Se1, n = 26). Left – Power split by median difference (Δ Se1-Se2) of accuracy of match; left middle – mis-match discrimination. Right middle – Power split by median z-score of neuropsychological assessment by CVLT total recall and right – learning slope. Note, the diverging steepness of the power spectrum above 30 Hz compared to the accuracy difference splits. Blue – better performance than median, red – worse performance than median.

